# Validating the use of bovine buccal sampling as a proxy for the rumen microbiota using a time course and random forest classification approach

**DOI:** 10.1101/2020.04.10.036665

**Authors:** Juliana Young, Joseph H. Skarlupka, Rafael Tassinari Resende, Amelie Fischer, Kenneth F. Kalscheur, Jennifer C. McClure, John B. Cole, Garret Suen, Derek M. Bickhart

## Abstract

Analysis of the cow microbiome, as well as host genetic influences on the establishment and colonization of the rumen microbiota, is critical for development of strategies to manipulate ruminal function toward more efficient and environmentally friendly milk production. To this end, the development and validation of noninvasive methods to sample the rumen microbiota at a large-scale is required. Here, we further optimized the analysis of buccal swab samples as a proxy for direct microbial samples of the rumen of dairy cows. To identify an optimal time for sampling, we collected buccal swab and rumen samples at six different time points relative to animal feeding. We then evaluated several biases in these samples using a machine learning classifier (random forest) to select taxa that discriminate between buccal swab and rumen samples. Differences in the Simpson’s diversity, Shannon’s evenness and Bray-Curtis dissimilarities between methods were significantly less apparent when sampling was performed prior to morning feeding (P<0.05), suggesting that this time point was optimal for representative sampling. In addition, the random forest classifier was able to accurately identify non-rumen taxa, including 10 oral and feed-associated taxa. Two highly prevalent (> 60%) taxa in buccal and rumen samples had significant variance in absolute abundance between sampling methods, but could be qualitatively assessed via regular buccal swab sampling. This work not only provides new insights into the oral community of ruminants, but further validates and refines buccal swabbing as a method to assess the rumen microbiota in large herds.

**IMPORTANCE:** The gastrointestinal tract of ruminants harbors a diverse microbial community that coevolved symbiotically with the host, influencing its nutrition, health and performance. While the influence of environmental factors on rumen microbes is well-documented, the process by which host genetics influences the establishment and colonization of the rumen microbiota still needs to be elucidated. This knowledge gap is due largely to our inability to easily sample the rumen microbiota. There are three common methods for rumen sampling but all of them present at least one disadvantage, including animal welfare, sample quality, labor, and scalability. The development and validation of non-invasive methods, such as buccal swabbing, for large-scale rumen sampling is needed to support studies that require large sample sizes to generate reliable results. The validation of buccal swabbing will also support the development of molecular tools for the early diagnosis of metabolic disorders associated with microbial changes in large herds.

## INTRODUCTION

The rumen is a specialized organ found in cattle that hosts a wide diversity of microorganisms from all three super kingdoms (for a review see (1, 2)). Essential to the digestion of complex plant polymers by the host, the rumen microbiota consists of several species of specialized fibrolytic bacteria capable of degrading lignocellulose (3). Microbial changes following total rumen exchanges (4) and some preliminary genome-wide association data (5, 6) suggest that the microbial community composition is unique to each individual cow and that the genetics of the host animal may influence community development/maintenance in the rumen. Unfortunately, the statistical determination of the extent of host-animal control over this phenomenon requires a large amount of input data and rumen microbial samples are often quite laborious to obtain.

Methods that directly sample the rumen contents of cattle are the rate-limiting step for generating a population-scale metric of the rumen microbiome. The gold-standard method for assessing rumen microbial contents is via rumen cannulation; however, this requires invasive surgery and cannot be performed on hundreds of cows in a herd. Stomach tubing is another method of sampling that provides direct access to rumen contents, but this method is labor-intensive and is uncomfortable for the cow (7, 8) (7). Given the requirements for surgery or labor-intensive sample collection, respectively, neither method is suitable for the development of a scalable industrial product. In light of the deficiencies of these methods, buccal swabbing has been proposed as a proxy for the rumen microbiota (9, 10). The ease of this method, combined with high-throughput sequencing of the 16S rRNA gene and its lower cost of implementation, make it a tantalizing option for obtaining population-scale rumen microbial samples.

Buccal swabbing is a noninvasive method that takes advantage of cattle rumination, an innate behavioral process that characterizes the ruminant clade of mammals (11, 12). During this process, the cow regurgitates, masticates, moistens, and swallows a bolus from the rumen, which is a mixture of previously ingested plant material that is resistant to prolonged chemical degradation. This process exposes additional surface area of the digesting plant matter to continued microbial fermentation (12). However, rumen microbes are not effaced from the surface of the bolus prior to mastication, and microbial DNA in the oral cavity may constitute a representative proxy of the rumen microbiota.

Indeed, the oral cavity has its own resident microbiota that contains both transient facultative anaerobes and feed-associated microbes (13, 14) that can be concurrently sampled during buccal swabbing. The identification and exclusion of these contaminants constitute a pre-requisite for the use of buccal swabs as proxy for the rumen microbiota (9). In previous studies, the depletion of these contaminants was performed with mathematical filtering based on the comparison of the relative abundances of a given taxa between rumen and buccal swab samples (9, 10). However, these approaches noted the need for further statistical and qualitative validation for wide-spread adoption of the technique due to confounding factors that could impact microbial taxa counts (9). This is a necessary step towards the use of buccal swabbing as an independent method, as future surveys may not always have access to paired rumen samples for calibration.

Previous surveys have also not considered sampling time as a potential confounding factor for interrogating rumen microbial counts via buccal swabbing (15–18). In the case of sample time, salivary dilution and contamination with feed silage communities could impact measured community composition and abundance. It is possible that there is a specific window of time in which buccal swab samples best mirror the rumen contents of the sampled cow. Prior to its widespread adoption as a suitable proxy for rumen sampling, buccal swabbing data must be compared in a modeling experiment to identify the magnitude of these biases.

In this study, we apply statistical learning methods to buccal swab data obtained from 21 cannulated Holstein cows to identify microbial taxa that are specific to the oral cavity. We hypothesize that the presence of non-rumen bacterial communities and the eventual salivary dilution of rumen microbial DNA impacts the comparability of buccal swab samples with in-situ rumen samples. We also tested if buccal swab OTU abundances can be used in regression models to determine the approximate abundance of rumen microbial genera in individual animals. Our analysis reveals an additional complexity in the diversity of microbes that colonize the ruminant gastrointestinal tract, and we expand the future use of buccal swabs in population-scale surveys of the rumen microbial community.

## MATERIAL AND METHODS

### Animal care and use

All animal procedures were conducted according to Research Animal Resource Center (RARC) protocol A005902-A02 approved on 07/28/2017 by the University of Wisconsin-Madison College of Agriculture and Life Sciences Institutional Animal Care and Use Committee. This work was carried out at the US Dairy Forage Research Center Farm, Prairie du Sac, WI, from 11/2017 to 06/2019 using a cohort of 21 cannulated lactating Holstein dairy cows (~2.5 years old) fed a total mixed ration in a free stall barn.

### Sampling

To identify the sampling time at which oral microbiota would best represent the rumen microbiota, paired oral (Buccal Swab, BS) and ruminal samples (Rumen Anterior Liquid, RAL; Rumen Anterior Solid, RAS; Rumen Ventral Liquid, RVL; Rumen Ventral Solid, RVS) were collected from 8 cannulated Holstein cows every 2 hours over the course of 10 hours, starting 1 hour prior to morning feeding (~ 9 AM) and ending just prior to evening feeding (~ 7 PM), totaling six time points (T1-T6). This dataset is hereafter referred to in the text as the summer time course (STC; see Table 1).

**Table 1.** Samples and experimental design.

| Sample set | Description | Sample count | Used in classification? | Used to train regression model? |
| --- | --- | --- | --- | --- |
| Summer, Time course, Farm 1 (STC) | Six timepoints of sampling paired buccal and rumen contents. | 8 animals | Yes | Yes |
| Spring sampling, Farm 1 (SPS) | Paired rumen and buccal contents; taken 4 hours after feeding | 5 animals | No | Yes |
| Summer sampling, Farm 2 (SUS) | Paired rumen and buccal contents; taken 2 hours prior to feeding | 8 animals | No | Yes |

Two other surveys of paired buccal swab and rumen content samplings were conducted on different animals in the same herd at two other timepoints separated by at least three months (Table 1). These datasets consist of a spring sampling (SPS; 5 cows) and a summer sampling (SUS; 8 cows) taken a year prior to the STC dataset. Swabs and rumen contents were processed in the same manner as listed for the time course survey, but samples were collected from animals four hours after feeding (all cows in SPS) or prior to feeding (all cows in SUS), representing equivalents to T4 and T1 from the time course trial, respectively. These samples were collected to provide additional power for training and testing regression models (see Table 1).

In all trials, two swabs (Puritan PurFlock Ultra sterile flocked swab with an 80 mm break point, Puritan Medical Products, Guilford, ME) were inserted in the buccal cavity of each cow and were gently scraped across the inner side of the right cheek for approximately 10 seconds. The buccal swabs were placed in a sterile conical tube (15 mL) containing 1 mL of sterile phosphobuffer saline and stored on ice during sampling. Immediately after buccal swabbing, rumen contents were collected via the rumen cannula and squeezed through double layers of cheesecloth to obtain an aliquot of 40mL of rumen liquids and 50 mL of a loosely packed rumen solid fraction. The solid fraction was squeezed once more to remove all liquids and the residual solid material was transferred to another container. All samples were stored and transported on wet ice and stored at −80 °C until processing and DNA extraction.

### DNA extraction and sequencing

Total genomic DNA was extracted from buccal swab, rumen liquid, and rumen solid samples as previously described (19). Sequencing was performed at the UW-Madison Biotechnology Center using the 2 × 250 bp paired-end method on an Illumina MiSeq following manufacturer’s guidelines (Illumina, Inc., San Diego, CA, USA). Detailed methods about and the library preparation and sequencing can be found in Skarlupka et al. (20).

### Bioinformatics analysis

DNA sequences were analyzed using mothur (v1.39.0) (21) as described previously (22). Coverage was assessed by Good’s index (23) and samples that displayed coverage less than 93% were discarded prior to normalization. To address differences in sequencing depths, the operational taxonomic unit (OTU) table was normalized by subsampling sequences to the sample with the smallest number of sequences and then normalizing across samples to produce equal sequence counts (3,000 sequences per sample). The normalized OTU table was used in further analyses as well as to calculate alpha diversity indices (i.e., Chao1 (24), Shannon (25), and Simpson (26)), Bray-Curtis dissimilarity index (27) as well as the relative abundance (reads/total reads in a sample x 100) of OTUs in each sample. Alpha diversity indices were calculated in mothur (v1.39.0) (21) whereas Bray-Curtis dissimilarity index was calculated using function vegdist available at R package vegan (v2.5-6) (28)

### Statistical analysis

All statistical analyses were performed in R (v3.6.1) and source code to reproduce these analyses is available in Supplementary Materials. Measurements of α-diversity (Chao1’s richness, Shannon’s evenness and Simpson’s diversity) and absolute abundance (i.e., sequence read counts) of OTUs detected in at least 80% of all samples, were assessed for normality and were found to follow a non-normal distribution. Differences in the alpha diversity indices and OTU absolute abundance values were analyzed, respectively, under Gamma and Poisson distributions, using a repeated-measure generalized linear mixed model estimated via penalized quasi-likelihood (29):

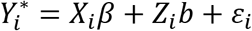

where 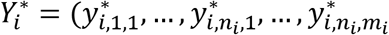 is a vector of Gamma- or Poisson-transformed of alpha diversity indices or OTU counts; *X_i_* is a design matrix relating individual observations to levels of fixed effects, *β* is a vector of fixed effects (i.e., sampling time, sample type, and their interaction), *Z_i_* is the incidence-matrix on random effects, *b* is the vector of random animal effects; *ε_i_* is a vector of random error terms. The resulting ANOVA P-values were adjusted for false discovery rate (FDR) using the Benjamini-Hochberg method, and values ≤0.05 were considered significant. Pairwise comparisons among the Least Squares Means (LSMEANS) were performed using Tukey’s Honest Significant Difference (Tukey HSD) method. In the presence of significant interaction effects, the LSMEANS of the sample types were compared within each sampling time. These analyses were performed using functions available at R package fitdistrplus (v1.0-14), MASS (v7.3-51.5), lsmeans (v2.30-0), and ggplot2 (v3.2.1) (30–33).

To visually explore the degree of dissimilarity between bacterial composition of oral and rumen samples collected at six distinct sampling times, Principal Coordinates Analysis (PCoA) was conducted on the Bray-Curtis distance matrix (27). In addition, Permutational Multivariate Analysis of Variance (PERMANOVA, nperm=1000) (34) with *post hoc* test using Benjamini-Hochberg correction was performed to assess differences in the composition of bacterial communities according to sample type, time points and their interaction. These analyses were performed using functions available in the R packages ggplot2 (v3.2.1), vegan (v2.5-6), and EcolUtils (v0.1) (28, 35, 36).

To identify taxa that discriminate between oral and rumen samples, a Random Forest classifier was trained on a random selection of 70% (162 samples) of the database composed of 232 samples and 2,031 OTUs and validated using the remaining 30% (70 samples). Only OTUs with relative abundance ≥ 0.05% present in at least one sample were included as input. The number of trees was set to 500, while the number of variables available for splitting at each tree node (mtry) was tuned and accuracy was used to select the optimal model using the largest value. In addition, to evaluate the capability of our model to predict on independent dataset, we adopted a repeated k-fold cross validation method (10-fold repeated 3 times). Prediction performance metrics (i.e., accuracy, sensitivity, specificity, precision and recall) and a confusion matrix were calculated and summarized by sample type. Finally, the Mean Decrease in Gini (i.e., Gini index) was used to calculate the variable importance score (VIMP) and select bacterial OTUs that were most predictive of sample types. To that end, we used the function varImp ((37)) that automatically scales the importance scores to be between 0 and 100. These results were plotted to show the most important sample type-associated bacterial OTUs with VIMP score ≥50%.

These analyses were performed using the R packages randomForest (v4.6-14) and caret (v6.0-85) (37, 38).

In order to evaluate if abundance of oral microbiota can be used to predict the abundance of rumen microbiota, we tested distinct regressions models (i.e., random forest, Random generalized linear model, GLMM zero-inflated quasi-Poisson). These analyses were performed using the R packages MASS (v7.3-51.5), caret (v6.0-85), randomForest (v4.6-14), and randomGLM (v1.02-1) (37–39).

### Data Availability

The raw sequence reads from all samples analyzed in this study are available on the NCBI Sequence Read Archive (https://www.ncbi.nlm.nih.gov/sra/) under the Bioproject accession number: PRJNA623113.

## RESULTS

### Amplicon sequencing and quality control

To provide metrics for quality control and optimal parameter selection, we sampled buccal and rumen contents from several cohorts of cannulated cattle (Table 1). To test if a difference in rumen sampling site had closer resemblance to swab samples, rumen strata (solids and liquids) from the anterior and ventral side of the rumen lumen were simultaneously collected. Samples are hereafter referred to by acronyms that denote their sample type (BS and R for buccal swab and rumen, respectively), and their location and content in the case of rumen samples (A, V, S, and L, for anterior, ventral, solid, and liquid, respectively). For example, the acronym RAL refers to a rumen anterior liquid sample. All samples were sequenced using the same methods and resulting data were processed using the same pipeline.

After sequence quality filtering and normalization, a total of 1,392,036 reads (mean 6,000.155 ± 132.615 SD per sample) and 196,258 OTUs (mean 845.94 ± 199.411 per sample) were obtained from 232 buccal, rumen solid, and rumen liquid samples in total. Good’s coverage estimation prior to normalization (0.969 ± 0.034 per sample) was deemed adequate and indicated that sequences sufficiently covered the diversity of the bacterial communities in our study. A full summary of sequencing statistics as well as rarefaction curves divided by sample type and time point is shown in Fig. S1 and Table S1.

Taxonomic composition analysis of the bacterial communities revealed a total of 2,031 OTUs (mean 112.46 ± 32.91 SD) present at relative abundances ≥0.05% and representing 20 phyla, 116 families and 279 genera. The average percentage of sequences unassigned to any phylum, family, or genus were 0.19 ± 0.15, 1.15 ± 0.45, and 10.49 ± 2.69, respectively. The most abundant OTUs, summarized at the phylum, family and genus levels according to sampling time and type are shown in Fig. S2.

### Time course analysis and sampling method comparability

We first sought to identify the effects of sampling method on the composition of observed microbial communities in the rumen. For this analysis, we used paired rumen strata (solid and liquid) and buccal swab samples taken from the STC cohort (see Table 1) in 2-hour intervals, with the first time point (T1) taken 1 hour prior to feeding. Rather than seeking a singular optimal time for sampling, we investigated the possibility that there are periods where the buccal microbial community may be less representative in terms of species prevalence and relative abundance of the rumen community.

Sampling type (i.e., buccal swabbing vs. rumen cannula sampling) had the largest effect on observed microbial content, as expected. Alpha diversity analysis revealed that Chao1 richness (number of species) varied significantly with sample type (P = 0.014) but not sampling time (P = 0.208) or the interaction of these two factors (P = 0.091). Shannon’s evenness (population density) and Simpson’s diversity (richness and abundance) varied with sample type (P < 0.001; P < 0.001), sampling time (P = 0.021; P = 0.047), and the interaction of these factors was significant (P < 0.001; P < 0.001). Regardless of sampling time, buccal swab samples displayed lower richness (i.e., Chao1) and evenness (i.e., Shannon), but higher diversity (i.e., Simpson) when compared to all types of rumen samples (Tukey HSD<0.05). Regardless of sample type, bacterial communities sampled at T3 and T4 displayed the lowest and highest Shannon’s evenness, respectively (Tukey HSD < 0.05). Significant differences in Shannon’s evenness and Simpson’s diversity were not observed between others timepoints (Tukey HSD < 0.05; Table S2). In regard to interaction terms, we observed that buccal swabs collected at T1 and T4 displayed similar evenness and diversity to all types of rumen samples. In contrast, buccal swab samples from other time points (T2, T3, T5 and T6) displayed lower evenness but higher diversity, relative to rumen samples (Tukey HSD < 0.05; Table S2).

We used PCoA to visually inspect the similarity of buccal swab samples to contemporary rumen cannula samples. In general, rumen samples grouped by phase (i.e., L vs S) rather than location (i.e., A vs V). Additionally, we found that bacterial communities from buccal swab samples obtained just prior to morning feeding (T1) grouped most closely to rumen solid samples (RAS + RVS) (Fig. 1). Moreover, ordination plots showed that T3 had the most pronounced differences between swab and rumen samples. The presence of higher OTU counts of silage-associated microbes belonging to the *Lactobacilli* in T3 suggest that feed contamination was a major contributor to this discrepancy (Figs. 2 and S2\).

**FIG 1.**
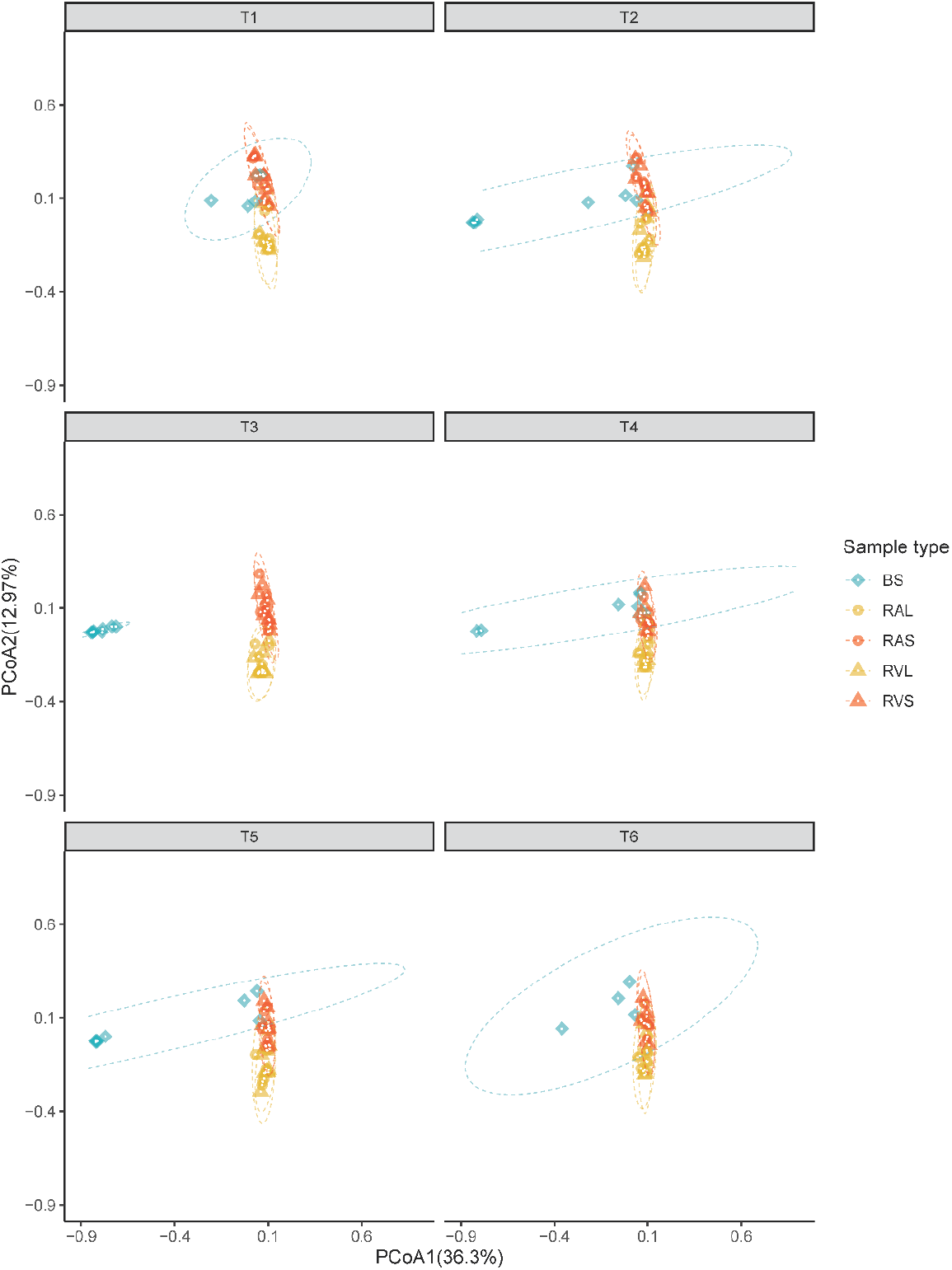
Principal coordinate analysis (PCoA) showing Bray-Curtis dissimilarities in the composition of bacterial communities between sample types within each sampling time. Individual points in each plot represent a dairy cow, different colors and shapes represent a sample type (BS: buccal swab, RAL: rumen anterior liquid, RAS: rumen anterior solid, RVL: rumen ventral liquid and RVS: rumen ventral solid), and each facet represents a time point (T1 to T6). Percentages showed along the axes represent, respectively, the proportion of dissimilarities captured by PCoA in 2D coordinate space.

**FIG 2.**
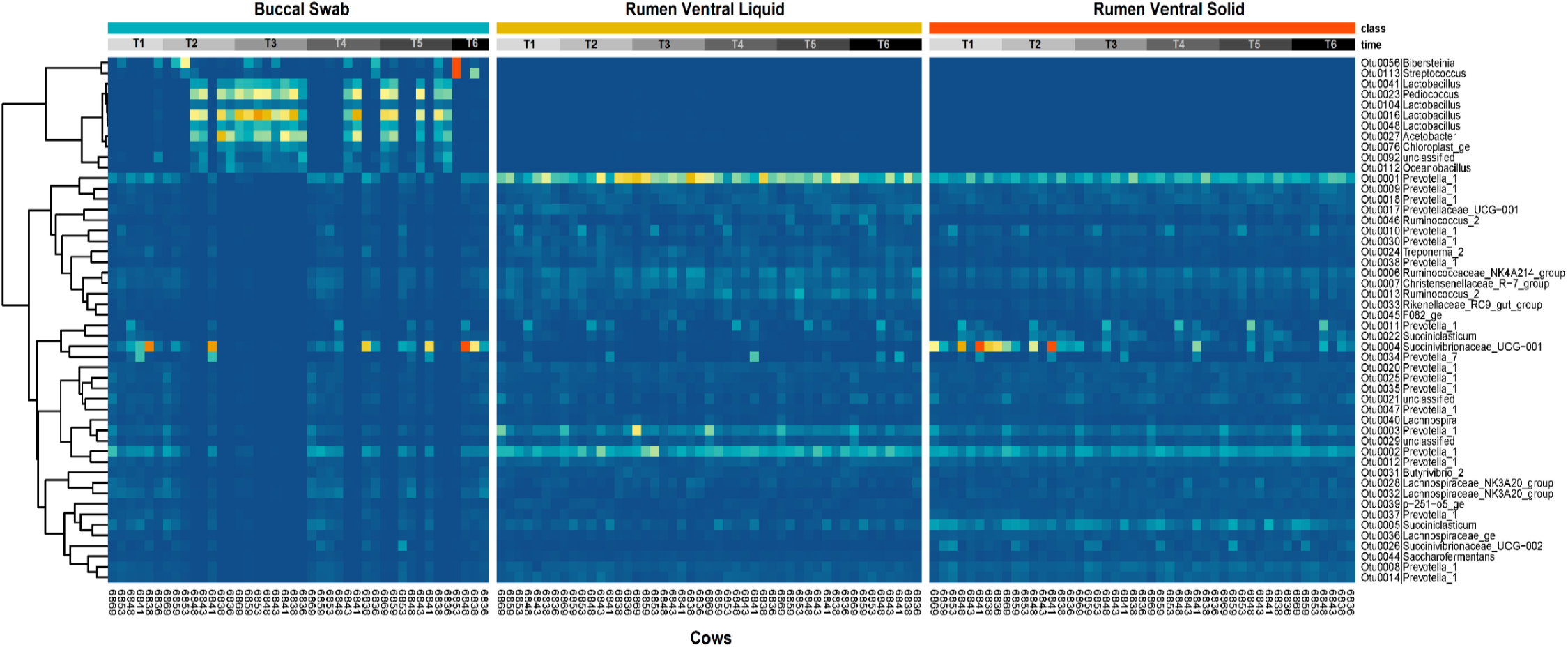
Distribution of the most abundant bacterial taxa among individual dairy cows according to sample type (BS: buccal swab, RVL: rumen ventral liquid and RVS: rumen ventral solid) and sampling time (T1:T6). The color-key represents the relative abundance at gradient of color from dark blue (low abundance) to dark orange (high abundance). The hierarchical dendrogram was established using Pearson product-moment correlations as the distance measure and “complete” as a clustering method.

**FIG 3.**
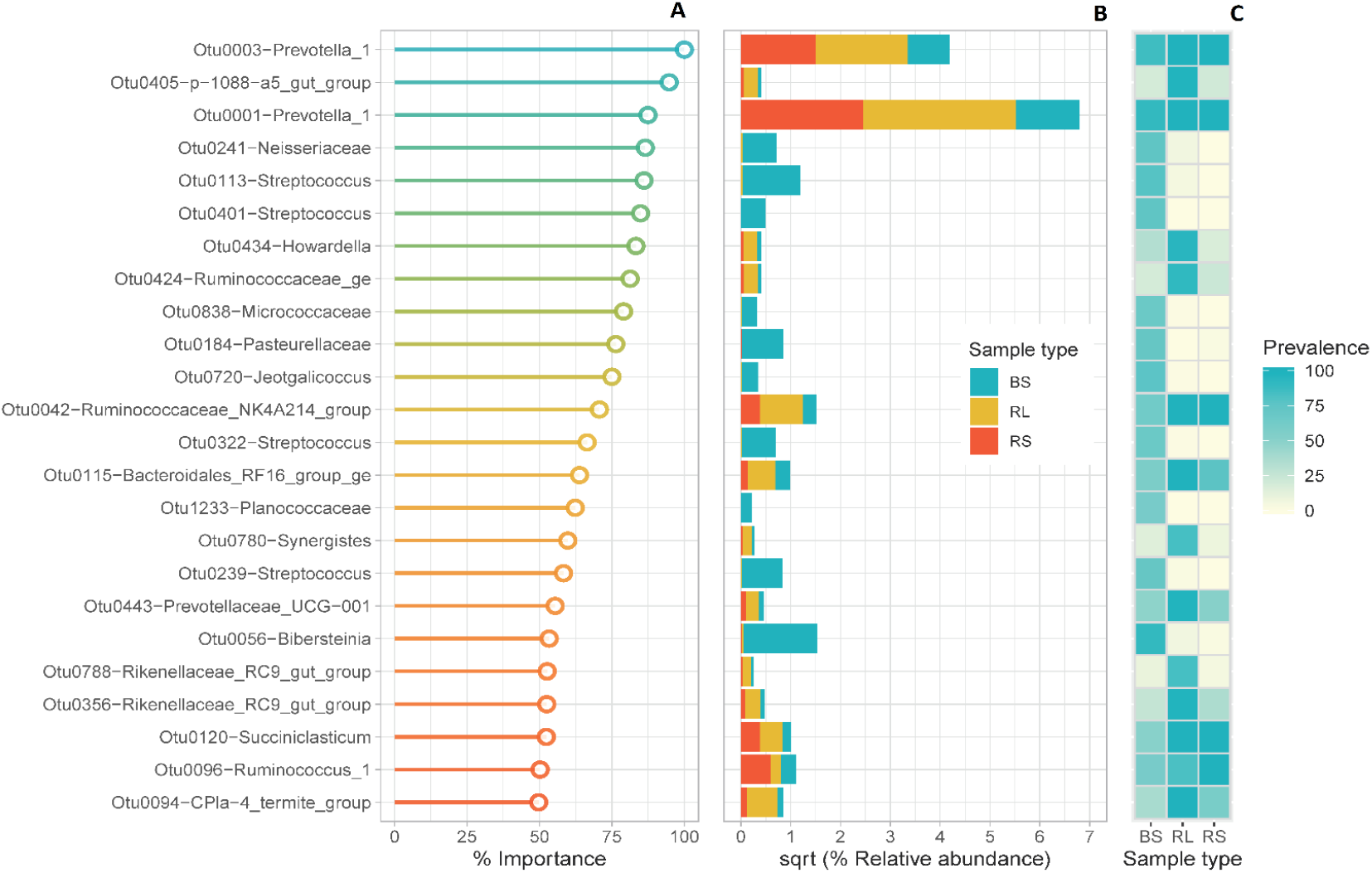
Variable importance (VIMP) plot from the random forest classifier. A) Lollipop chart showing the most important bacterial signatures that displayed importance (% Mean Decrease in Gini≥50) and that discriminate between buccal swab (BS), rumen liquids (RL) and rumen solids (RS) samples. B) Bar-plots of sqrt-relative abundance of OTUs according to sample type; C) Heat map of prevalence of OTUs in each sample type.

**FIG 4.**
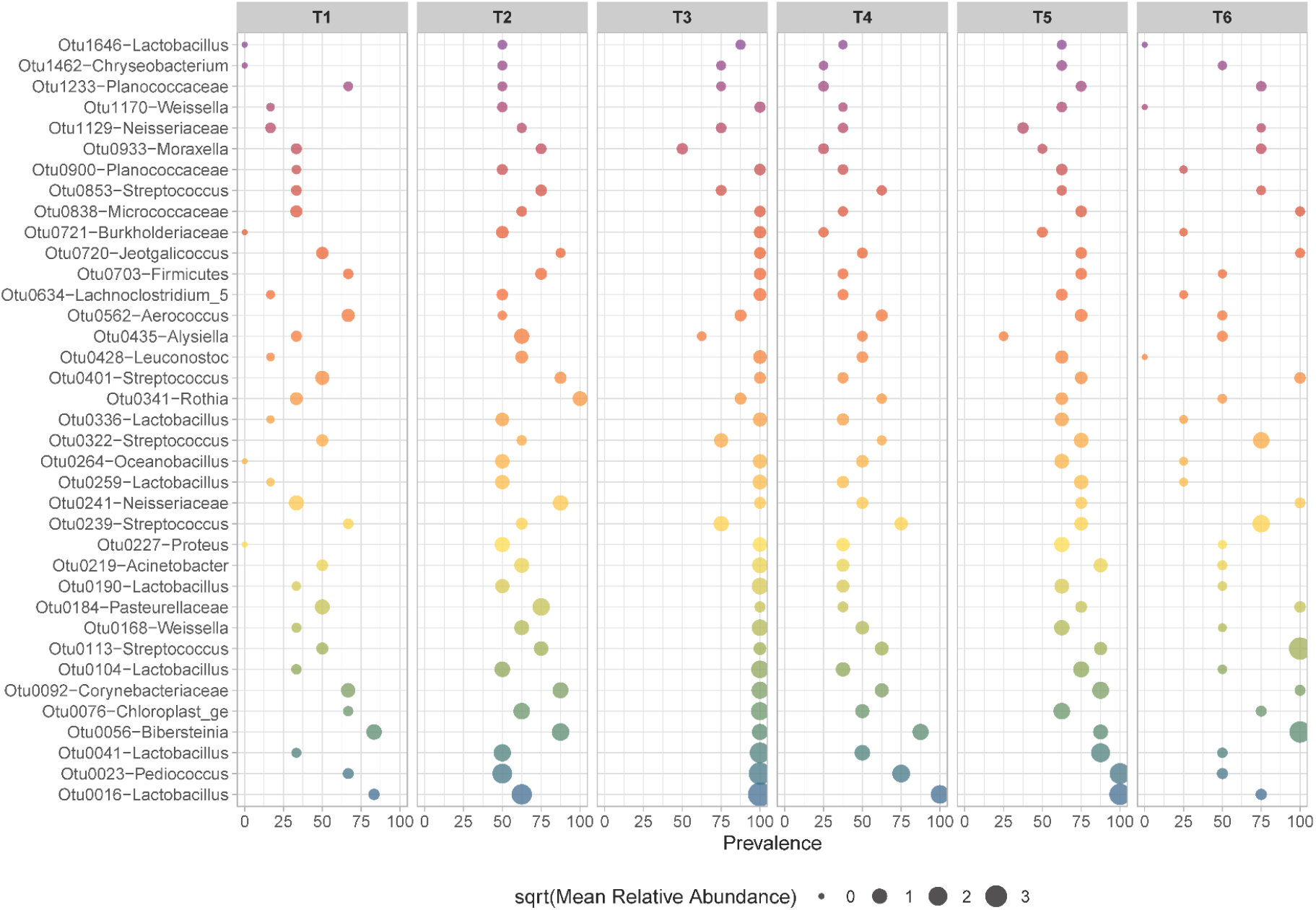
Bubble chart showing the prevalence and relative abundance of the oral OTUs assigned to higher taxa (phylum, family or genus level) according to sampling time (T1:T6).

PERMANOVA showed that Bray-Curtis dissimilarities in the composition of bacterial communities were significantly driven by sampling time (R squared= 0.044, P< 0.001), sample type (R squared= 0.284, P< 0.001), as well as by the interaction of these two factors (R squared= 0.106, P< 0.001). Pairwise comparisons between sample types showed that the composition of BS samples differs significantly from all types of rumen samples (P=0.010). In addition, we found that bacterial composition at sampling time T1 was significantly different from T3 (P = 0.015) and T5 (P = 0.045). Lastly, comparisons between sample types within each sampling time indicates that the composition of bacterial communities in BS samples is similar to those observed in the RAS samples only at T1 (P = 0.054), confirming the clustering observed in the PCoA (Fig.1 and Table S3).

In addition to compositional dissimilarity, we assessed differences in the absolute abundance (i.e., read counts) of 277 bacterial OTUs (prevalence of at least 80% of all samples) in response to sampling time, sample type and the interaction of these two factors (Figs. 5, 6, and Table S7). Overall, most of the variance in the absolute abundance of bacterial communities in our study was ascribed to interaction terms given that 240 OTUs varied simultaneously with sampling time and sample type. Meanwhile, the differences ascribed to main effects were far less apparent, given the abundance of only 38 and 20 OTUs that varied independently in response to sample type and sampling time, respectively (Table S7).

**FIG 5.**
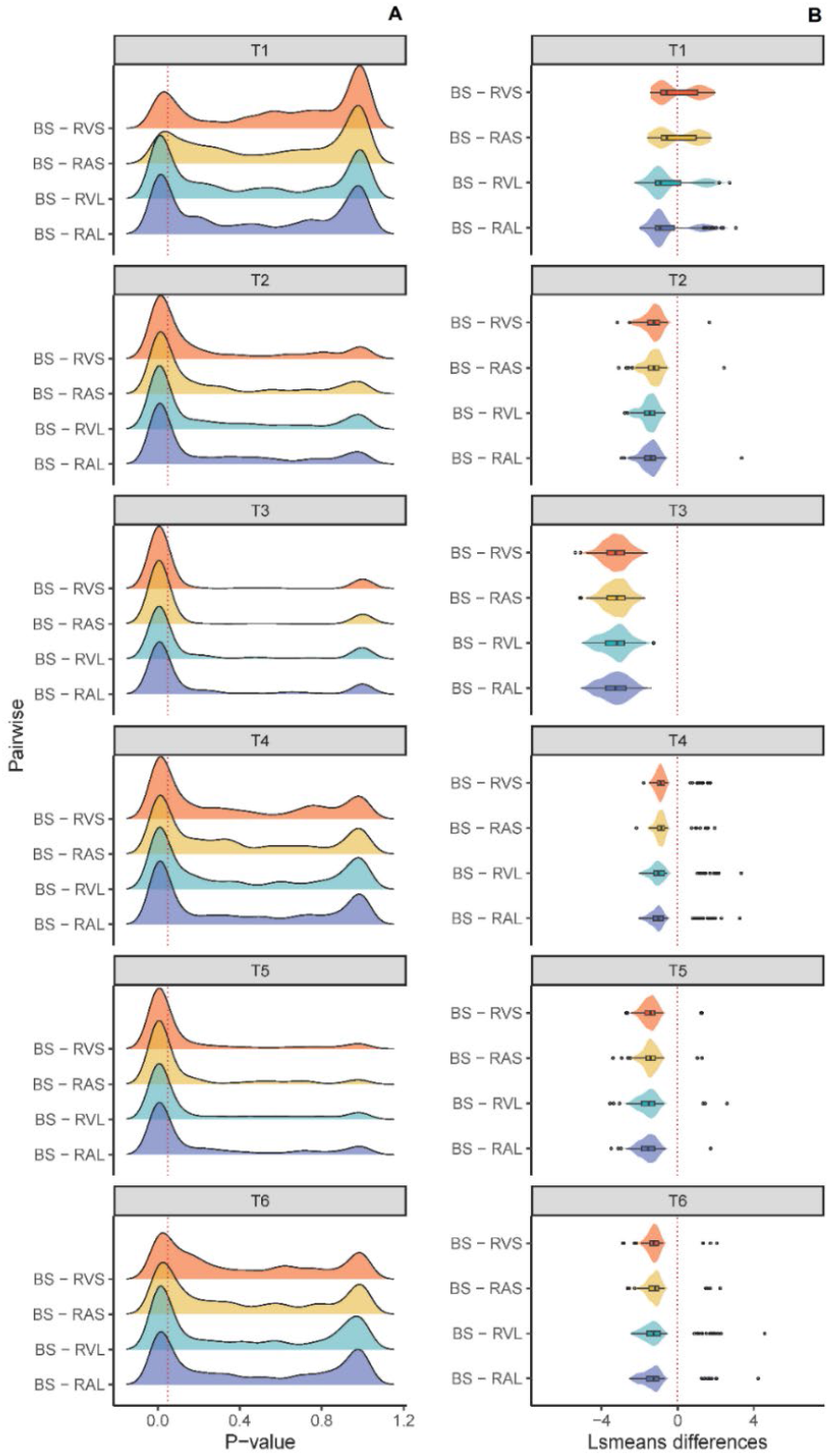
**A)** Ridgeline plots showing the distribution of bacterial OTUs whose abundance varied significantly (red line = P-value ≤ 0.05) in pairwise comparisons between buccal swab (BS) and all types of rumen samples (RAL, RVL, RAS, and RVS) within each sampling time (T1:T6). **B)** Violin plot showing the Least Squares Means (LSmeans) differences of significant pairwise comparisons (Tukey HSD ≤ 0.05) between buccal swab and all types of rumen samples within each sampling time.

**FIG 6.**
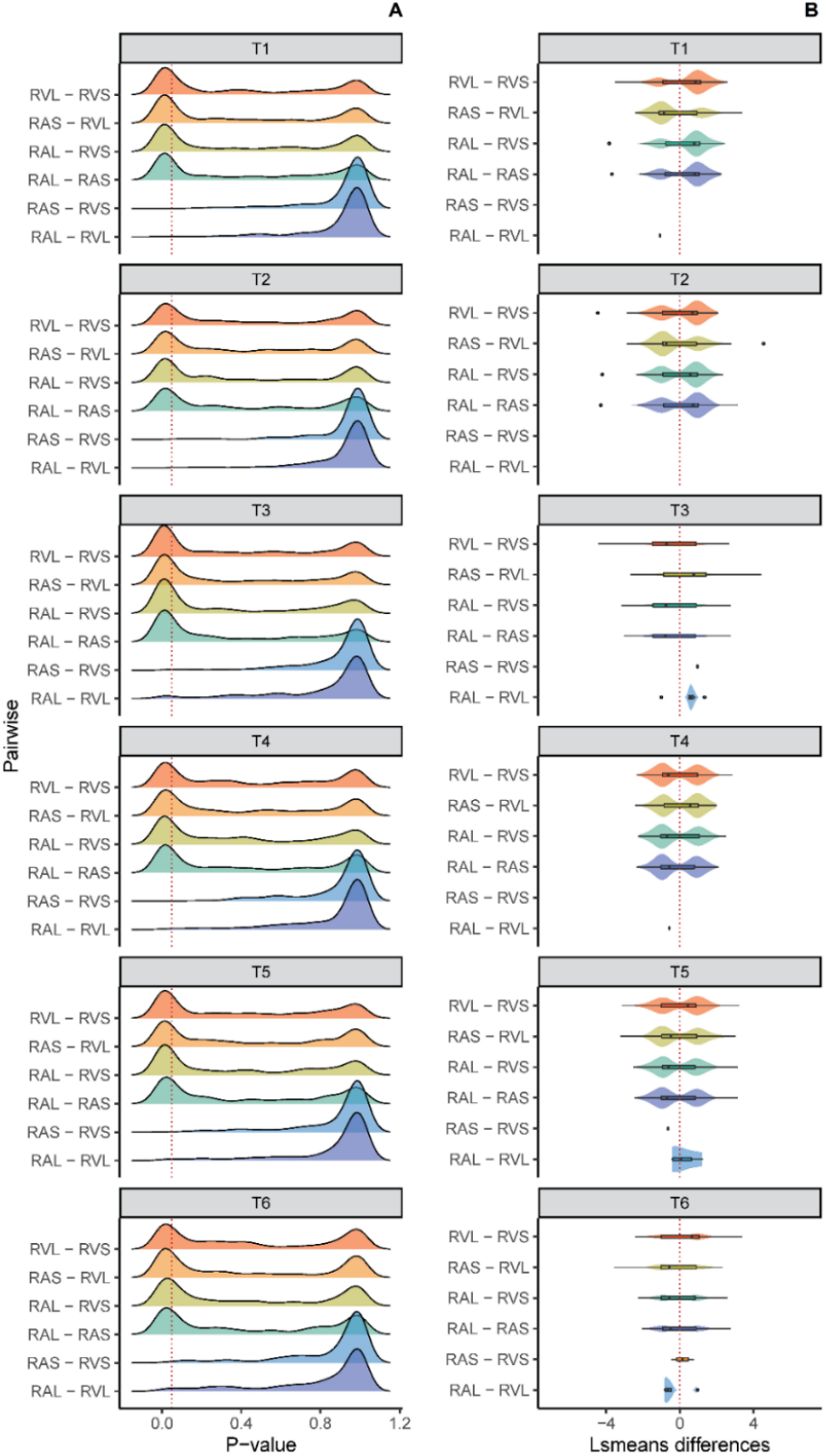
**A)** Ridgeline plot showing the distribution of bacterial OTUs whose abundance varied significantly (red line=P-value≤0.05) in pairwise comparisons between all types of rumen samples (RAL, RVL, RAS, and RVS) within each sampling time (T1:T6). **B)** Violin plot showing the Least Square Means (LSMEANS) differences of significant pairwise comparisons (Tukey HSD ≤0.05) between all types of rumen samples within each sampling time.

Comparisons between sample types within each sampling time showed that fewer OTUs had significantly different absolute abundance between buccal and rumen samples taken at T1 followed by T4 and T6 (Fig. 5A). At these particular time points, the significant differences in the absolute abundance of OTUs between buccal swab and rumen samples were less pronounced than observed at other sampling times (Tukey HSD ≤ 0.05; Fig. 5B). In contrast, greater significant differences in the absolute abundance of OTUs between BS and all rumen samples were observed at T3 followed by T5 and T2 (Figs. 5A, 5B, and Table S8).

In addition, no significant differences in the absolute abundance of OTUs between RAS and RVS were observed at T1, T2 and T4. However, some OTUs varied in absolute abundance between RAL and RVL at others sampling times, mainly at T3 followed by T5 and T6 (Fig. 6A and Table S8). Pronounced differences in the absolute abundance of several OTUs between liquids and solids contents were observed at all time points. Specifically, the majority of the OTUs sampled at T1 and T2 displayed higher absolute abundance in rumen liquids than in rumen solids (i.e., RAL vs. RAS and RVL vs. RVS) while the opposite was observed at other time points (Fig. 6A, 6B, and Table S7).

Regardless of sample type, comparisons performed between sampling times showed that the absolute abundance of bacterial OTUs were significantly lower at T3 and T5 in comparison to the other time points, particularly with T4 and T1 (Figure S3 and Table S8). Finally, comparisons performed between sample types showed that absolute abundance of bacterial OTUs were significantly lower in buccal swabs than all types of rumen samples (Tukey HSD ≤ 0.05), regardless of sampling time. These differences were less apparent when buccal swab and rumen solids were compared (see Figure S3 and Table S8). However, a few exceptions were observed for OTUs assigned to Prevotellaceae_Ga6A1_group and Succinivibrionaceae_UCG-002, whose absolute abundance were significantly higher in BS in comparison to rumen liquids (RAL or RVL; Tukey HSD < 0.05) (Table S8).

### Random forest classifier analysis

We next sought to identify key microbial taxa present in the oral microbial community that contributed to discrepancies observed in our ordination plots. To statistically distinguish between taxa that had differences in relative abundance in each sample type, we trained a random forest classifier model using the STC cohort samples. Random forest is a supervised learning algorithm which uses ensemble learning method (i.e., combine several trees base algorithms) to construct better predictive performance (for a review see (38, 40) and has been widely and successfully employed for classification and regression purposes. In a classification problem, the algorithm returns a list of predictor variables (i.e., bacterial OTUs) that can be ranked according to their individual importance (i.e., VIMP score) in classifying the data.

Our preliminary analyses showed that the overall performance of the random forest classifier using five classification categories for sample type (BS, RAL, RAS, RVL, and RVS) was quite low (Accuracy 58.6% and Kappa 48.2%), even after estimation and tuning of model hyper-parameters (Table S4). This result supports the observation of high similarity between bacterial communities from rumen solids (RAS and RVS) and liquids samples (RAL and RVL) from different rumen lumen areas as observed in the PCoA (Fig. 1). We found improved classifier accuracy when rumen samples were merged based on rumen content strata (liquids and solids) into a single type in the training and testing sets (collectively referred to as RL and RS, respectively). This merger unbalanced our training set by providing a two-fold increase in rumen categories (RL and RS = 95 samples each), and we thus implemented a re-sampling method for future model training to prevent misclassification of our minority class (BS = 42 samples). We tested three additional re-sampling methods (i.e., under-sampling, over-sampling, and Synthetic Minority Over-sampling Technique, SMOTE) to prevent classification bias towards the majority classes (41, 42). The results showed that random forest trained with additional re-sampling using the SMOTE had higher performance metrics than the other methods (Table S5).

Our final model was able to predict sample type-associated bacterial features with high accuracy (97.78% ± 3.7%) and Cohen’s kappa values (96.3% ± 5.4%). Cohen’s kappa is a frequently used statistic to assess the performance of machine learning models under a multi-class classification problem and or unbalanced data (43, 44). Other performance metrics such as sensitivity, specificity, precision and recall were also calculated for each sample type and are presented in Table S5. Additionally, our classifier returned the variable importance score (VIMP), as a function of the Mean Decrease in Gini, of each bacterial OTU, which can be used to discriminate between oral and rumen samples (Table S6). Thus, higher values of VIMP score expressed as a percentage indicate higher feature importance (i.e., bacterial OTU) in discerning between classes and, in our case, between sample types.

### OTU categorization based on variable importance estimates

Bacterial OTUs with high VIMP scores (≥ 50% mean decrease Gini) displayed patterns that allowed for manual categorization. Based on average taxon prevalence per sample type and sampling time, we categorized these OTUs into three categories: core, oral, and rumen (Table 2, Fig. 3 and see Table S6 in the supplementary material). The remaining OTUs whose VIMP score was lower than 50% were also categorized for the sake of completeness but were not analyzed further (Table S6). The core category consisted of OTUs that displayed moderate to high prevalence (≥60 to 100%) in all sample types (both rumen and buccal) consistently across timepoints. The rumen category was defined as the community well represented (prevalence ≥75%) in rumen liquids and/or solids, and was underrepresented in buccal swab samples (prevalence<60%) at all time points (Fig. 3, Table 2 and Table S6). Finally, the oral group consisted of OTUs well represented in buccal swab samples (prevalence≥60%) but were either absent or underrepresented in the rumen samples (<60% prevalence) across time points. The oral group was found to contain silage community microbes (i.e., *Lactobacilli*) at time points where feed was provided to the animals (e.g., T3, see Fig. 4), further supporting our classification and the model’s accuracy.

**Table 2.**
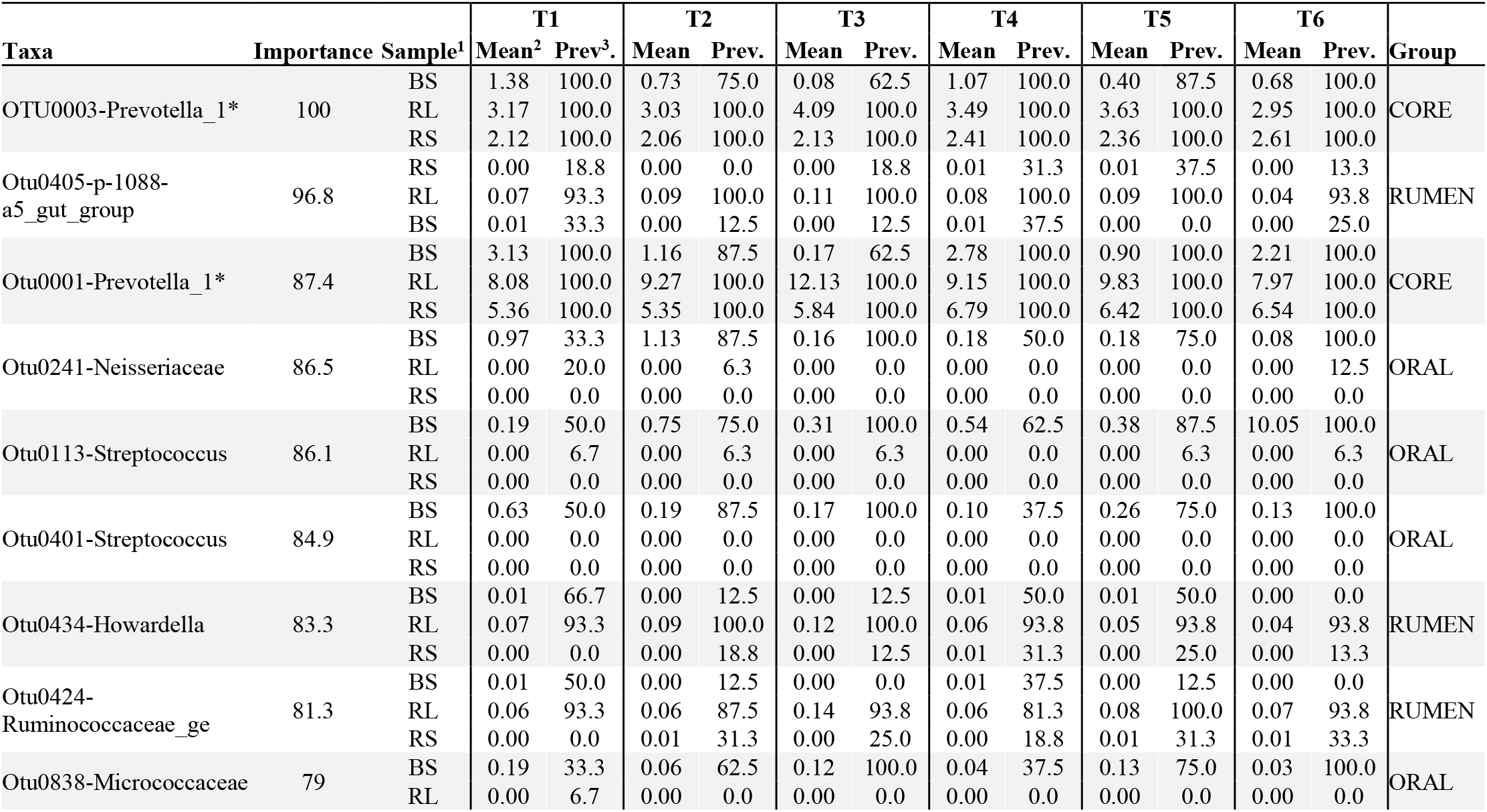
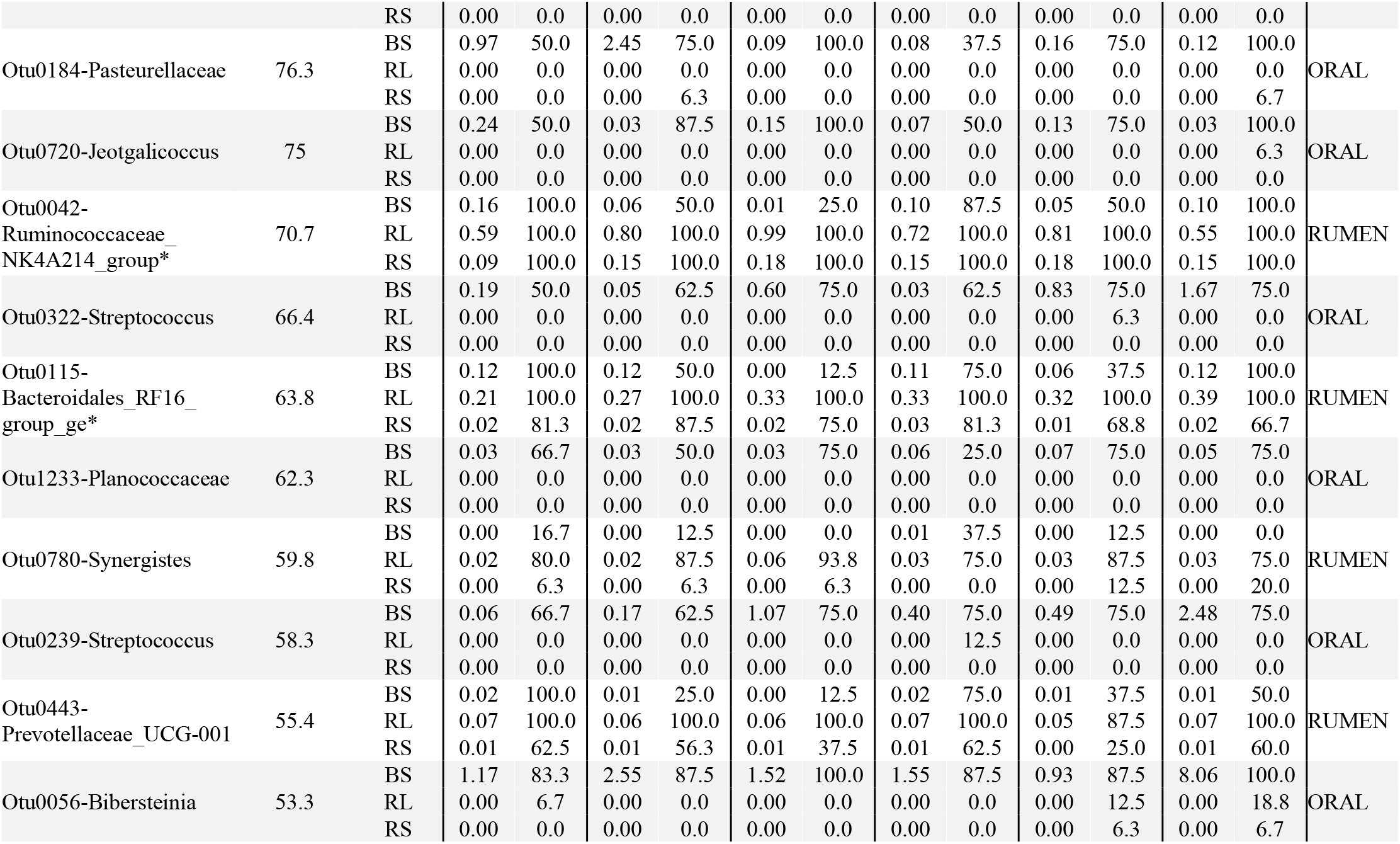
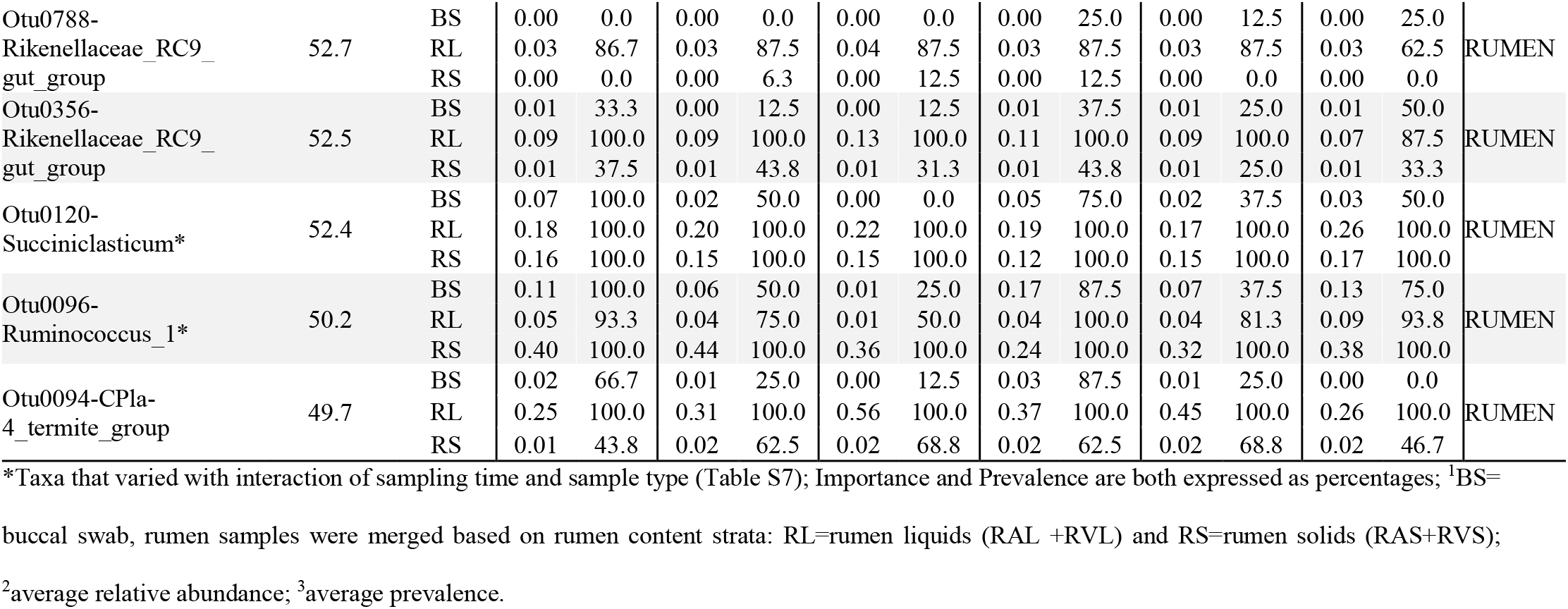
Variable importance analysis from the random forest classifier showing the most important bacterial OTUs (importance: scaled Mean Decrease in Gini≥50%) that discriminate between buccal swab and rumen samples.

In the core group, we identified two OTUs (VIMP>80%) assigned to the genus *Prevotella_1* (Fig. 3 and Table 2) that displayed high prevalence in both buccal swab and rumen (liquid and solid) samples. The absolute abundance of these taxa was significantly lower (Tukey HSD≤0.05) in buccal swabs than in rumen samples (Tables S6 and S7). This suggests that these taxa can be reliably sampled via swabbing but that their absolute abundances are greatly biased compared to the paired rumen samples.

We also identified taxa in the families Neisseriaceae, Pasteurellacea, Micrococcaceae, and Planococcacea, as well as in the genera *Streptococcus, Jeotgalicoccus*, and *Bibersteinia*, which displayed moderate to high VIMP scores (≥ 50%) and were assigned to the oral category. These taxa were overrepresented in terms of prevalence and abundance in buccal swab samples and displayed very low or zero abundance in rumen liquid and solid samples (Fig. 3 and Tables 2 and S6). In addition, we observed that several oral taxa (i.e., *Oceanobacillus, Lactobacilli, Lachonoclostrium*, *Leuconostoc*, *Rothia*, and *Proteus*) were underrepresented in terms of abundance and prevalence at specific time points, including T1, T4 and T6, relative to time points T2, T3 and T5 (Fig. 4 and Table S6).

Finally, the classifier also selected rumen strata OTUs that have lower relative abundance in the buccal swab samples (rumen category). Several were specific to rumen liquids (0405-p-1088-a5_gut_group, *Howardella*, Ruminococcaceaa_ge, *Synergistes*, Prevotellaceae_UCG-001, Rikenellaceae_RC9_gut_group) and others were derived from the rumen solids (*Ruminoccocus_*1, Prevotellaceae_UCG-001 and *Oribacterium*) whose overall importance was ≥33% (Fig. 3 and Tables 2 and S6).

### Random forest regression analysis

We next sought to test whether the abundance of OTUs found in buccal swab samples could be used to predict the abundance of rumen OTUs. We tested the ability of four linear models (random forest regression, three log-linear models with either a Poisson distribution, zero inflated, or random generalized linear model (RGLM)) to characterize the relationship between bacterial OTUs of paired buccal swab and rumen liquid samples. In order to provide additional data for our training regression models, we incorporated data from 21 cows sampled in two other surveys (Table 1) processed with the same methods used for the time course study. It is important to note that random forest regression was performed using sequence relative abundances whereas log-linear models use sequence absolute abundance (i.e., number of reads) for each OTU, assuming a Poisson distribution of read counts. Our random forest and Poisson regression model converged, but exhibited low accuracy in cross-validation studies as shown by a low coefficient of determination (R-Squared = 0.39 ± 0.05) and high Root Mean Square Error (RMSE = 0.28 ± 0.09). We attempted to tune additional parameters in the random forest model, but were unable to achieve an accuracy R-Squared above of 0.42 ± 0.07 on a per-OTU basis. Conversely, zero inflated and RGLM trials failed to converge, despite several attempts to filter the OTU tables and tune model parameters. These results may be related to our use of a small dataset as well as the non-linear relationship between the buccal swab and rumen OTU abundance/counts on a per-sample basis.

## DISCUSSION

In this study we evaluated the ability of the buccal swabbing method to describe bacterial communities found in two types of rumen samples taken at six distinct sampling times over the course of ten hours. Buccal swab samples are an attractive alternative to more labor-intensive methods of sampling the rumen microbial community, but may suffer from bias due to contamination by the surrounding oral community (9, 10). We first sought to identify the effect of sampling time on buccal swab community composition as we hypothesized that animal rumination patterns and salivary flow may change the relative abundance of key members of the rumen community.

Our time course analysis suggests that there is a small, but statistically significant, effect of sampling time on the comparisons of several buccal swab microbial taxa with contemporary rumen samples from the same animal. After dividing sampling times into two-hour intervals, we sampled buccal contents from each animal just prior to the start of morning feeding (T1), within regular intervals during and after feeding (T2, T3, T4, and T5), and prior to evening feeding (T6). We found that the only major outlier was at time point 3 (T3), where the greatest dissimilarities in the bacterial communities between buccal swabs and rumen samples were observed. It is possible that additional contamination by the silage microbial community and increased salivary flow induced by feeding changed the relative abundance of key rumen taxa in the oral samples of cows sampled at T3. This is evidenced by the presence of *Lactobacilli* from silage communities in the buccal swabs, but not in the rumen contents (Fig. 2 and 4). Our results support a hypothesis that there are brief windows of time in which buccal swab data best represent contemporary rumen microbial data. This means that future surveys will need to record time of sampling relative to animal feeding in order to standardize results.

We also tested the possibility that buccal swab samples may be compositionally similar to rumen content fractions taken from different positions in the rumen (i.e., Anterior vs Ventral).

Our comparisons of sampling time and sample types found no differences between the bacterial communities of the anterior and ventral rumen microbial communities, which prevented us from finding such an association (Fig. 1). This result is likely associated with the constant mixing of rumen contents due to the contractions of the reticulorumen, which would result in indistinguishable variation in our observed rumen microbial OTU counts (12). This finding contrasts from previously published work that identified noticeable differences in sample composition from five different locations of the rumen lumen via PCR DGGE surveys (45). We therefore cannot rule out the possibility that our sampling and analysis methods could not identify the small effects that these locations have on the community.

We also found greater similarity between bacterial taxa present in buccal swabs and rumen solids than in rumen liquids (Fig. 1). We suspect that this reflects a key stage of the rumination process whereby, immediately after regurgitation, the liquid fraction of the bolus is swallowed (12). It is possible that the bacterial taxa that are predominant in the liquid-phase of the rumen contents are evacuated from the oral cavity early in the process of rumination. During mastication of the bolus, bacteria from solid-phase of rumen contents are more likely to adhere to oral mucosal surfaces and are more likely to be sampled during buccal swabbing.

In order to identify non-rumen taxa in buccal swab samples, we employed a machine learning classifier to assist in the filtering of oral and silage microbial communities in buccal swab samples. As has been noted previously (9), the presence of the commensal oral microbial community in buccal swab samples prevents direct comparisons between rumen content samples and buccal swabs and must be filtered from buccal swab samples prior to analysis using manual and mathematical methods (9, 10). By using a random forest classifier, we were able to assign importance estimates to individual microbial taxa based on their use as a feature in our classification models, as has been done previously (46, 47). The top OTUs, after variable importance analysis, consisted of microbes that were oral-specific (oral, n = 10), rumen-biased (rumen, n = 12), and those with high prevalence regardless of sample type but varied based on relative abundance (core, n = 2). These findings support our observations of the influence of sample type on OTU relative abundance, and also identified members of the oral-microbial community that were prevalent only in buccal swab samples. In addition, the top OTUs identified by our VIMP analysis included two members of the *Prevotella*, which were found to vary substantially between buccal and rumen samples (Table S7). These two OTUs were prevalent in all samples and at all time points; however, their relative abundance in buccal swabs was lower than in the rumen samples. These differences were far less apparent at T1, which as just prior to feeding, than at any other sampling time. This observation of similarity at only one time point implies that sampling time had a large effect on the estimated relative abundance of this clade, as confirmed by our ANOVA.

The OTUs present within the oral category represent taxa that are poorly represented in buccal swab samples. Indeed, we identified commensal oral microbes from the genus *Rothia* that were present only in the buccal swab samples (the oral category). These taxa can be safely removed from future buccal swab surveys. We also identified several oral taxa (i.e., *Lactobacillus*, Chryseobacterium, *Burkholderiaceae*, *Oceanobacillus*) that were prevalent at some time points, and underrepresented or even absent at others (Fig. 4) showing that sampling time is a critical factor to be considered in future studies. The higher prevalence of these taxa during (T2) and immediately after (T3) feeding suggests that these sampling times will result in buccal swab data that is least representative of the rumen contents of the animal.

Our use of random forest classifiers suggests that machine-learning methods can be used to approximate the rumen microbial community at the time of sampling. More accurate estimation of these communities will be beneficial to rumen microbial ecology experiments that suffer from low sample counts. However, we were unable to achieve an acceptable rate of error (measured via residual error of observed and predicted OTU counts) from our regression analysis. We found that multicollinearity of predictors and weak linear association between oral and rumen OTUs prevented accurate regression. We suspect that other factors (i.e., sampling time, herd, diet) must be controlled for in the modelling of these data, as evidenced by significance of sampling time and interaction terms in our PERMANOVA and ANOVA. Moreover, it is possible that the taxonomic affiliation of our OTU counts could be masking individual species level abundances that provide far more variance than expected for the regression model. Similarly, our genus-level assignments could also contain inaccuracies due to strain abundance differences in the oral cavity vs. the rumen contents.

Finally, we cannot rule out the possibility that several OTUs are metabolically active (i.e., facultative aerobes) in both locations and can proliferate in the oral cavity, thereby creating a non-linear relationship between their abundance estimates in buccal swabs and rumen contents. While this presents an impediment to the use of buccal swabs for classical microbial ecology experiments, we note that buccal swab data is still useful for other associative analysis. The ability to collect large numbers of samples from a diverse cohort of animals can present an opportunity for associations of microbial profiles with animal production and performance metrics including milk production, health and even fertility phenotypes. Such experiments would benefit from the removal of biases that we identified in this survey.

In summary, we have identified significant effects of sampling time and sample type on the composition of rumen microbial OTU counts derived from buccal swabs and rumen samples. The buccal swab samples were prone to significant bias based on the time of sampling, with specific time points showing higher prevalence of the oral- or feed-associated microbial community than others. For future surveys using buccal swabs as a proxy for rumen microbial counts, we recommend buccal sampling at least 2 hours prior or four hours after feeding. Our data also suggests that a portion of the rumen microbial community will remain inaccessible to buccal swab samples; however, this bias may not necessarily impede future association studies with host animal phenotypic traits.

## ACKNOWLEDGMENTS

This research was funded by a USDA Agricultural Research Service (Washington, DC) CRIS project 5090-31000-026-00-D supporting D.M.B., J.C.M, and J.Y, a USDA ARS CRIS project 5090-31000-025-00D supporting K.F.K, a USDA ARS CRIS project 8042-31000-001-00-D supporting D.M.B., and a USDA ARS CRIS project 8042-31000-002-00-D supporting J.B.C. This work was also supported in part by a USDA National Institute of Food and Agriculture (NIFA), Agricultural and Food Research Initiative (AFRI) Foundation Grant #2019-05592 to G.S., D.M.B., and J.B.C. J.Y. was also partially supported by a USDA NIFA AFRI Foundation Grant #2015-67015-22970. J.H.S. was supported by NIH National Research Service Award T32 GM07215.

Mention of trade names or commercial products in this article is solely for the purpose of providing specific information and does not imply recommendation or endorsement by the USDA. The USDA is an equal opportunity provider and employer.

We would like to thank Paul J. Weimer for conversations related to the design of this study, and for his careful reading and suggestions for this manuscript. The authors declare no conflicts of interest.

